# Upregulation of the Proto-Oncogene Src Kinase in Alzheimer’s Disease: From Molecular Interactions to Therapeutic Potential

**DOI:** 10.1101/2024.02.07.579336

**Authors:** Diego Mastroeni, Chun Kit Chan, Nader Morshed, David Diouf, Camila de Ávila, Crystal Suazo, Jennifer Nolz, Ulia Lopatin, Qi Wang, Geidy Serrano, Thomas Beach, Travis Dunkley, Kendall Jensen, Daniel Van Den Hove, Forest M. White, Po-Lin Chiu, Abhishek Singharoy, Eric M. Reiman, Benjamin P. Readhead

## Abstract

Alzheimer’s disease (AD) is a progressive neurodegenerative disease, resulting in an irreversible deterioration of multiple brain regions associated with cognitive dysfunction. Phosphorylation of the microtubule-associated protein, Tau, is known to occur decades before symptomatic AD. The Src family of tyrosine kinases are known to phosphorylate select tyrosine sites on Tau and promote microtubule disassembly and subsequent neurofibrillary tangle (NFT) formation. Our data show that the proto-oncogene, non-receptor tyrosine kinase Src colocalizes with a range of late (PHF1) to early (MC1) AD-associated phosphorylated Tau epitopes. The strongest co-occurrence is seen with MC1 (probability of MC1 given Src =100%), an early AD-specific conformational dependent epitope. Single-cell RNA sequencing data of 101 subjects show that *Src* is upregulated in both AD inhibitory and excitatory neurons. The most significantly affected, by orders of magnitude, were excitatory neurons which are the most prone to pathological Tau accumulation. We measured Src phosphorylation by mass spectrometry across a cohort of 48 patient neocortical tissues and found that Src has increased phosphorylation on Ser75, Tyr187, and Tyr440 in AD, showing that Src kinase undergoes distinct phosphorylation alterations in AD. Through Brownian dynamics simulations of Src and Tau, we show that as Tau undergoes the transition into disease-associated paired helical filaments, there is a notable seven-fold increase in Src contact with Tau. These results collectively emphasize Src kinase’s central role in Tau phosphorylation and its close association with Tau epitopes, presenting a promising target for potential therapeutic intervention.

## Introduction

Alzheimer’s disease (AD) is the most common form of dementia, a highly debilitating disorder affecting more than 40 million people across the world^1,2^. AD is a progressive neurodegenerative disease, resulting in an irreversible deterioration of multiple brain regions associated with cognitive dysfuntion^1,3^. Alzheimer’s disease encompasses a continuum; it is now recognized that pathophysiological changes begin decades before clinical manifestations. The spectrum of AD spans from clinically asymptomatic to severely impaired, but neuropathological evidence of disease can be evident by the third decade of life ^4^.

Characteristic neuropathological features of AD were first described more than 100 years ago: extracellular amyloid plaques (Aβ-plaques), and the intra-neuronal accumulation of an abnormal form of the protein Tau (neurofibrillary tangles (NFT))^3^. However, these hallmarks can coexist with several physiological and morphological changes in the brain such as chronic brain inflammation, cerebral amyloid angiopathy, microglial activation, astrogliosis, and dystrophic neurites^1,2,5^. Taken together, these processes drive neurotoxicity, neuronal loss, mitochondrial dysfunction, synaptic dysfunction, and ultimately brain atrophy ^5^. However, the origin and underlying mechanisms through which these physiological changes lead to the pathophysiology of AD are not yet known. For this reason, predictions of the rate of preclinical changes and the onset of the clinical phase are extremely difficult to formulate, complicating the design and timing of therapeutic interventions aimed at modifying the course of the disease.

Recent evidence has pointed to a class of proteins whose actions are altered in AD, and whose disruption might be explanatory for several AD features. The Src kinase family belongs to the class of non-receptor tyrosine kinase^6^. This family includes several members, whose main components are: Src, Yes, Fyn, Lck, Lyn, Frk, Fgr, and Blk ^6-9^. Under normal conditions, Src family kinases interact with numerous cytosolic, nuclear, and membrane proteins by phosphorylating tyrosine residues and modulating intracellular signaling ^10-13^. *In vitro* findings show a potential role of Src-family kinases in the development of several AD-associated hallmarks ^14-18^. Indeed, Aβ-oligomers are a strong activator of intracellular non-receptor tyrosine kinases ^14-19^. The over-activation of this family of kinases leads to the disruption of several fundamental neuronal processes such as synaptic transmission, regulation of cytotoxicity, apoptosis, and re-entry into the cell cycle ^19-23^. Given the multiple avenues by which Src kinases interact with Tau and its various phosphorylation states, alterations in *Src* expression could lead to abnormal activity and the formation of full-blown NFTs. Here we show that *Src* is upregulated in multiple AD brain regions, highly expressed in excitatory neurons, colocalizes with the earliest disease-associated Tau epitopes, and binds paired helical filaments (PHF).

## Methods

### Clinical and Pathological Assessment for histochemical and Western blot studies

Subjects were all volunteers in the Arizona Study of Aging and Neurodegenerative Disorders (AZSAND), a longitudinal clinicopathological study of aging, cognition, and movement in the elderly since 1996 in Sun City, Arizona. Autopsies are performed by the Banner Sun Health Research Institute Brain and Body Donation Program (BBDP; www.brainandbodydonationprogram.org).^24^ For control subjects, no memory complaints or history of memory complaints, cognitive function within 1.5 standard deviations of the age- and education-adjusted norms, CeradNP-Not AD, MMSE score 30-29 (inclusive), Braak staging II, and NIA-AA-No AD. For AD cases, the following criteria were applied to ensure a clear distinction from control subjects: presence of persistent and escalating clinical memory complaints, cognitive function exhibiting a decline of more than 1.5 standard deviations from age- and education-adjusted norms, CeradNP-probable to definite AD, Mini-Mental State Examination (MMSE) score below 12, and Braak stage IV, NIA-AA intermediate and high for AD, indicative of the pathological progression associated with AD. Early disease states were chosen (e.g., Braak stage IV) to ensure the full spectrum of disease-associated Tau conformation and phosphorylation changes. Later Braak stages (Braak stage VI) show very high level of ghost tangles^25^ which minimizes the number of neurons that may be reactive for some of the earlier disease states (e.g., MC1). All subjects sign Institutional Review Board-approved informed consents allowing both clinical assessments during life and several options for brain and/or bodily organ donation after death.

### DAB Immunohistochemistry

Immunohistochemical studies were completed on 28 human hippocampal tissues (BA27), superior frontal gyrus (BA11), middle temporal gyrus (BA21), entorhinal cortex (BA28) and substantia nigra pars compacta. For each region, the same 10 AD, 9 NC, and 9 PD samples were used. AD, ND, and PD samples were selected with the following criteria: For AD, samples were devoid of other clinical and neuropathological neurological diseases, Braak stage IV, CeradNP-probable to definite AD, MMSE of 11 to 9, plaque density moderate to frequent, and NIA-AA intermediate and high for AD. ND samples were devoid of clinical and pathological changes associated with neurological diseases besides those that are described as findings consistent with normal aging. All ND samples were Braak stage II, CeradNP-not AD, MMSE of 30 to 29, plaque density zero to sparse, and NIA-AA not AD. For PD subjects, samples were devoid of clinical and pathological changes associated with neurological diseases besides those associated with PD. All PD samples were Braak stage II-III, CeradNP-not AD, MMSE of 30 to 27, plaque density zero to sparse, NIA-AA not AD, Unified LB stage III. Brainstem/Limbic. All samples were matched for PMI, age, and sex, among other covariates (see table of samples, **Supplementary Table 1)**. For detailed methods, please see^26-28^. Briefly, 40um free-floating sections were washed in PBST and blocked in H2O2, then 3% bovine serum albumin (BSA). Following blocking, steps tissues were incubated in primary antibodies (**Supplementary Table 2**), overnight at 4°C. Sections were washed and then incubated in species-specific secondary (1:1000, Vector) for two hours at room temperature. Sections were washed incubated in 1:1000 avidin/biotin reagent, washed, and incubated in DAB. All sections were reacted simultaneously, dried, taken through graded alcohols, cleared in xylene, and mounted using permount. Deletion of primary antibody or incubation with blocking peptide resulted in the abolition of specific immunoreactivity in all cases. Adjacent serial sections were stained with cresyl violet for cell layer identification and verification that the CA1 of the hippocampus was intact.

### Double label Immunohistochemistry

For fluorescence microscopy, the sections were washed 3X in PBST, blocked with either 3% normal goat serum or 3% BSA, and incubated for 2h. After further washing, sections were incubated in primary antibodies overnight, washed again, and incubated in species-specific, fluorophore-conjugated secondary antibodies (**Supplementary Table 2**). After a final wash, the sections were mounted, taken through Sudan Black to reduce autofluorescence, and coverslipped with Vectashield mounting media (Vector). All sections were counterstained with 4’,6’-diamidino-2-phenylindole (DAPI) (Thermo Fisher) before mounting.

### Immunohistochemical Statistics

Slides were imaged using Olympus IX71 or Nikon A1R HD25 confocal imaging system. Immunoreactivity was analyzed using Image J software. To provide estimates of Src immunoreactivity, bright field intensity (DAB) / fluorescence intensity analysis was performed using Image J software (Image J, U.S. National Institutes of Health, Bethesda, MD; imagej.nih.gov/ij/). Intensity measurements were corrected for background differences by dividing the measured intensities by the average intensity of a background region in each section. For colocalization studies, 2 fields at 40x magnification were counted per subject (∼40 neurons/subject). Colocalization (co-occurrence) or how many pixels in a segmented image are overlapping between the red (594nm) and green (488nm) channels. The acquired images undergo segmentation, a process where regions of interest are identified through pixel intensity values. To quantify the degree of overlap between signals from different channels, colocalization analysis is performed. This involves applying thresholds to pixel intensity values to create binary masks for each channel, highlighting areas with significant signal. Colocalization was calculated to measure the proportion of signal overlap between the red and green channels.

### Western blot

As previously described in detail^28^, proteins were isolated and separated by electrophoresis. The adapted protein quantity per lane was loaded into 4-20% 1.5 mm Tris-glycine 15 well mini gels (Bio-Rad) with 1X Laemmli sample buffer. Gels were run in running buffer, containing Tris Base, Glycine, and 10% SDS, at a constant 100 V for 60 minutes. Proteins were then transferred to nitrocellulose (Bio-Rad) and immersed into transfer buffer (10X CAPS, Methanol, and dH2O), at a constant 100V for 1 hour. Membranes were blocked using 5% BSA and 0.05% Tween in 1X PBS at room temperature for 1 hour and then probed with the primary antibody (anti-Src, Mouse monoclonal, 1:2000 in 5% dry milk) overnight at 4ºC with gentle shaking. Membranes were then washed and incubated with the secondary antibody dilution (Horse anti-mouse, HRP-ligated, PI 2000, 1:5000 in 5%dry milk, Thermo Fisher) for 1 hour at room temperature, with gently shaking. After being exposed to chemiluminescence substrate (Pierce, Supersignal West Pico), membranes were imaged on Amersham Imager 680 detection system and analyzed using ImageJ software. Each Blot was then stripped and re-probed for B-actin (loading control). The Optical density (OD) measures of each band were taken by using a sampling window of constant size. Groups were compared by using Analysis of variances (ANOVA) tests, and regression analysis, with a 95% confidence interval, and significance at α = 0.05.

### snRNA-seq data analysis: Clinical and Pathological Assessment for SnRNA-seq

Subjects were all volunteers in the Arizona Study of Aging and Neurodegenerative Disorders (AZSAND), described above. Most subjects are clinically characterized with annual standardized test batteries consisting of general neurological, cognitive and movement disorders components, including the Mini Mental State Examination (MMSE). Subjects for the current study (n=101) were chosen by searching the BBDP database for a full spectrum of AD neuropathology, in the absence of other neurodegenerative disease diagnoses. The complete neuropathological examination was performed using standard AZSAND methods.^24^ The neuropathological examination was performed in a standardized manner and consisted of gross and microscopic observations, the latter including assessment of frontal, parietal, temporal and occipital lobes, all major diencephalic nuclei and major subdivisions of the brainstem, cerebellum and spinal cord (the lattermost only for those with whole-body autopsy). Detailed clinical data, postmortem neuropathological data, and demographics of the cohort are described in.^29^ We constructed gene co-expression and detected multiscale gene modules using MEGENA^30^on the differentially abundant excitatory neurons, which were identified as those neurons in control subjects most susceptible to neuronal loss in AD. Differential expression of genes between AD vs control was identified by the FinderMarkers function of the Seurat (v4.0) workflow^31^, using the MAST algorithm^32^. *NEUROD2* and *Src* coexpression correlations in excitatory neuron subtypes were calculated from the snRNA-seq data in^33^ by a pseudo-bulk approach, using the sum of the gene counts from all the nuclei within the subclusters normalized by the voom function from the R package limma^34^, taking sex, age at death, education and PMI as covariates. Correlation coeffients were calculated by the lm function in R.

### Mass Spectrometry: Clinical and Pathological Assessment for mass spectrometry studies

Subjects were all volunteers in the Arizona Study of Aging and Neurodegenerative Disorders (AZSAND) described above. Samples of human middle temporal gyrus (MTG) were secured from AD, or neurologically normal, non-demented (ND) elderly control brains. Cognitive status of all cases was evaluated antemortem by board-certified neurologists, and postmortem examination by a board-certified neuropathologist resulting in a consensus diagnosis using standard NIH AD Center criteria for AD or ND. The AD and ND groups were well matched for age, sex, and PMI. ND control cases: ND Braak I n=5, ND 74 Braak IV n=18 (23 total); Age: 84.1 ± 7.1 years; Sex: 12 females and 11 males; Postmortem interval 75 (PMI: 2 hours 58 minutes +/- 75 min. AD cases: AD Braak IV n=10, AD Braak V n=8, AD Braak VI n=7; Age: 76 84.3 +/- 7.9 years; Gender: 11 females and 14 males; PMI: 3 hours 23 min +/- 180 min. For additional sample data see.^35^ To analyze the AD proteome and phosphoproteome, we generated proteolytic peptide digests of middle temporal gyrus (MTG) cortical brain tissue from patients with late-onset AD and age-matched, non-diseased (ND) controls. Batches of AD and ND peptide samples were multiplexed with TMT10-plex. TMT-labeled peptides were immunoprecipitated with anti-phosphotyrosine antibodies (4G10, PT66) followed by Fe^3+^ IMAC phosphopeptide enrichment. Enriched peptides were loaded onto a C18 column and analyzed by LC-MS/MS using a 140 min gradient. Ions were dynamically selected for fragmentation using a top-20 untargeted method. Peptide identification and quantification were performed using Proteome Discoverer and MASCOT. Peptide abundances were normalized by dividing by the mode relative abundance in each sample. Peptide abundances across TMT10-plex batches were normalized using the average of all ND samples without reported amyloid pathology. Tau cluster centroid values were taken directly from the original publication describing this dataset^36^.

### Brownian Dynamics

Computational studies on Src-Tau interaction were performed in a X-step manner. First, representative structures derived from experiments were chosen as starting models for simulating Src and Tau. For Src, PDB 1YOJ, which represented an active Src^37^, was employed. Compared with the inactive form of Src, active Src is more flexible and allows more protein residues to be solvent accessible. This enhanced exposure facilitates our simulation to sample Src residues that are susceptible to associate with Tau. For Tau, 3 different structures were employed. The first structure was constructed using Protein Data Bank (PDB)^38^, representing Tau aggregating on a microtubule of healthy individuals. The second and the third models were from PDB 6NWP and PDB 6NWQ, which, respectively, represented the cores of the type I and type II Tau filament of Chronic traumatic encephalopathy (CTE) and AD.^39,40^ Folds in CTE Tau, are structurally identical to those in AD,^41^ and are denoted as paired helical filaments (PHFs) and straight filaments (SFs). These structural models of Src and Tau were converted to respective mean field representations suitable for Atomic-Resolved Brownian Dynamics (ARBD), following our previous work^42^. Lastly, we launched 400 replicas of 4-microseoncd-long ARBD simulations for each of the following cases: (1) Src diffusing around Tau microtubule; (2) Src diffusing around Tau disease model I; and (3) Src diffusing around Tau disease model II. These simulations were then all converted back to molecular dynamic trajectories to analyze Src-Tau associations and Src diffusion pathways around different Tau aggregates.

## Results

### Src is upregulated in multiple AD brain regions

To determine if Src protein levels were upregulated in one or multiple brain regions we analyzed the substantia nigra pars compacta (SN), entorhinal cortex (BA28), hippocampus (BA27), middle temporal gyrus (BA21) and superior frontal gyrus (BA11) in Normal Control (NC), Parkinson’s disease (PD) and AD subjects (**Fig.1**). PD and SN (the pathogenic region associated most clearly with PD) were used to determine whether deviations in Src are a general feature of neurodegeneration or are AD-specific. In the SN, all three conditions show pigmented nigral neurons, but only in AD cases did we observe positive Src immunoreactivity (IR) (**Fig. 1C insert**), indicating that the upregulation of Src is not a generalized neurodegenerative phenomenon. All three, NC, PD, and AD showed cortical and hippocampal Src IR (**Fig. 1D-L**). It is to be expected in an aging population to have some neurofibrillary tangle (NFT)-like reactivity, even in PD tissues^43^. Comparison of AD, PD, and NC however, showed clear separation (**Fig.1**). Src staining in AD samples revealed intense neuropil and neuronal IR, reminiscent of NFT labeling in our previous studies^27^. The strongest neuropil staining was in the entorhinal cortex (**Fig. 1F**), and the most significant neuronal IR was observed in the hippocampus (**Fig. 1I**) followed by the middle temporal gyrus (**Fig. 1L**). The neuropil is comprised of mostly dendrites, axons, and synapses^44^ and thus provides a proxy measure of total neuronal connectivity. The EC is heavily interconnected with the neocortex and hippocampus, and is one of the earliest, if not the earliest brain region affected^45^, indicating that dysregulation of Src in the EC may substantially precede dysregulation in other less reactive brain regions. These results show that Src is highly expressed in multiple brain regions in AD and is not generally observed in other neurodegenerative diseases like PD.

**Figure 1:**
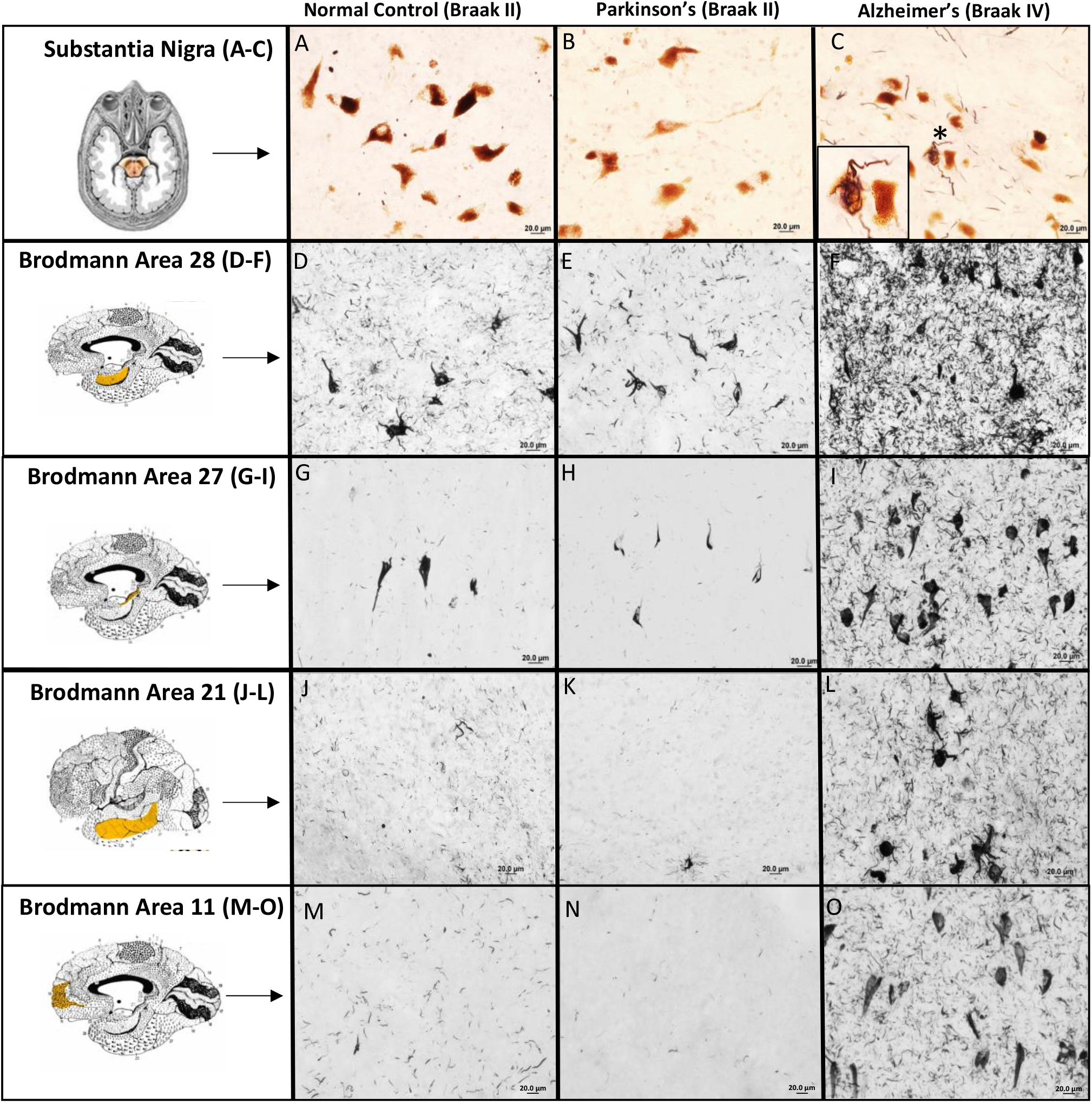
Representative Src immunoreactivity (IR) in four brain regions: substantia nigra pars compacta (SN), entorhinal cortex (BA28), hippocampus (BA27), middle temporal gyrus (BA21) and superior frontal gyrus (BA11) in Normal Control (NC), Parkinson’s (PD) and Alzheimer’s (AD) subjects. In the SN only AD cases showed positive IR, see insert (C). All three, NC, PD and AD showed cortical and hippocampal IR. More Src IR was observed in AD entorhinal cortex, hippocampus, and middle temporal gyrus, compared to NC and PD subjects. The most intense neuropil immunoreactivity in AD subjects was observed in BA28. PD cases were included as well as the PD-related pathognomonic region (SN) to show AD specificity. Note, neurons in the SN are naturally pigmented (brown).

To quantify Src protein levels we performed western blots of NC (Braak II/III) and AD cases (Braak IV (**Fig. 2**). Braak IV AD cases were used to evaluate the earliest disease state according to Braak^25^. Our results show a significant (p<.005) upregulation of Src protein in AD middle temporal gyrus, consistent with our histochemical study (**Fig. 1J vs 1L**).

**Figure 2:**
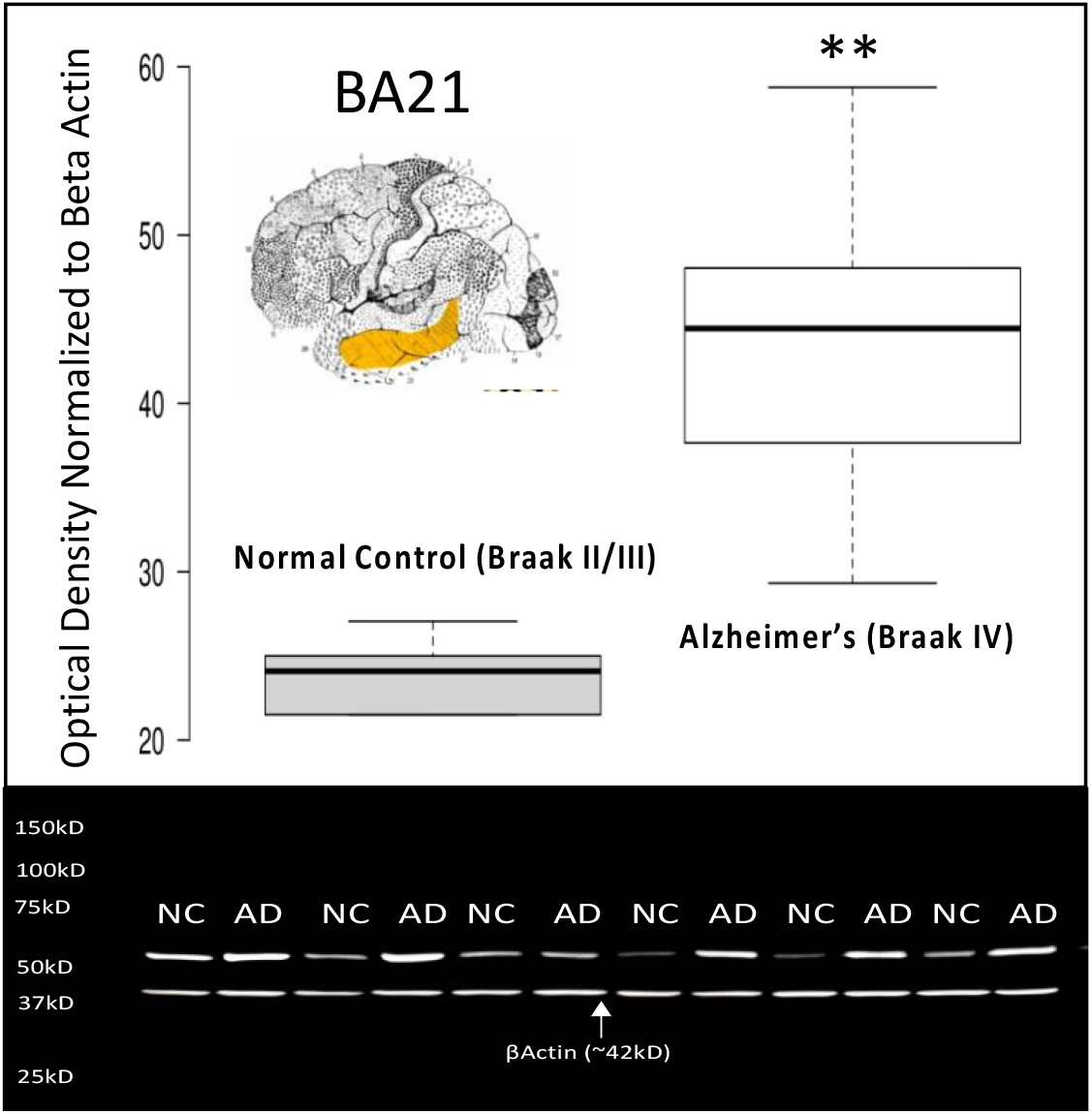
Western blot of Src protein levels in age-matched control and AD subjects. Complementary to IHC results we observed an overall increase in SRC expression in AD MTG (BA21). **p<.005. Western blot was normalized (protein loading) using beta actin.

### Src colocalizes with multiple disease-associated phosphorylated Tau epitopes

To determine whether the tangle-like IR in Figure 1 was associated with known markers of NFTs we performed colocalization studies using a commercial Tau antibody P-tau23, an end-stage phospho-Tau marker PHF-1, and early Tau markers PG5, CP13, and MC1 (**Fig. 3**).^46^ For our colocalization studies, we specifically examined the CA1 region of the hippocampus. This choice was informed by substantial evidence pointing to early tangle formation and the progression of disease in this area.^47,48^ In the context of degenerating nerve cells, PHF-1 or ps396/ps404 represents an end-stage disease state.^49^ In contrast, MC1 (a conformation-dependent antibody targeting the epitope within aa 312-322) is recognized as the earliest disease state.^46,49^ The other three antibodies used in the study—PG5 (which identifies PKA-dependent phosphorylation of ser409), CP13 (phosphorylated at serine-202), and P-tau231—are considered intermediate disease states. Notably, all five Tau antibodies were found to colocalize with Src, as depicted in Figure 3. The strongest co-occurrence with Src was with early Tau marker MC1 **(Fig. 3A**, probability of MC1 given Src =100%). The weakest co-occurrence with Src was with the late Tau marker PHF1 (**Fig. 3E**, probability of PHF1 given Src =43%). Intermediate molecules P-tau231(**Fig. 3B**) CP13 (**Fig. 3C**), and PG5 (**Fig. 3D**) showed intermediate probability given Src 61%, 58% and 54% respectively (**Supplementary Table 3**). Collectively, this data shows that Src is most strongly associated with early disease states.

**Figure 3:**
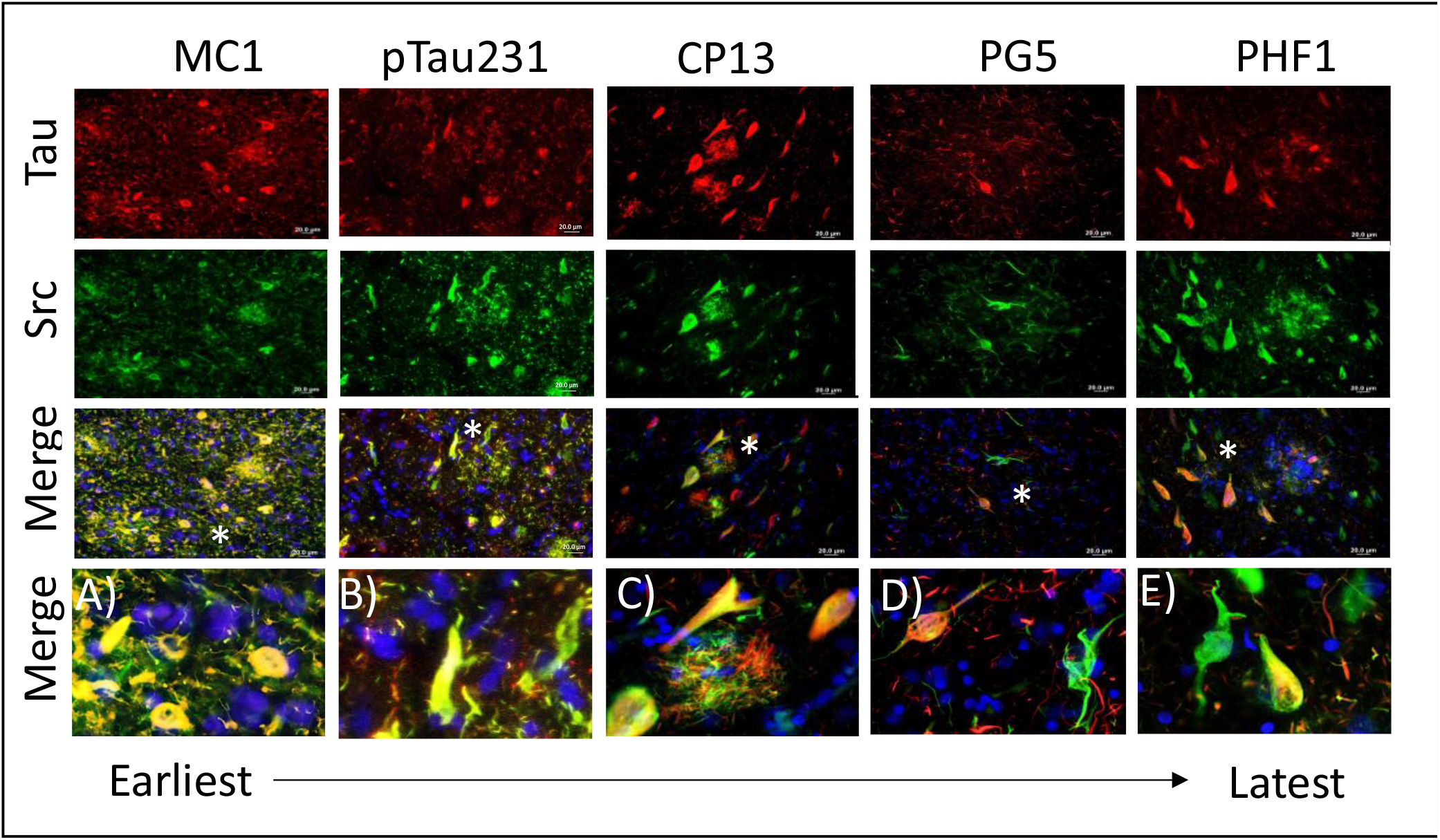
Representative double labeled photomicrographs of Src (green) and select early tau markers MC1, P-tau231, CP13, PG5 and late-stage P-tau marker PHF-1 and (red). The strongest association with Src was with early tau marker MC1 (A) and weakest, PHF1 (E). Sections were counter stained with DAPI.

**Figure 4:**
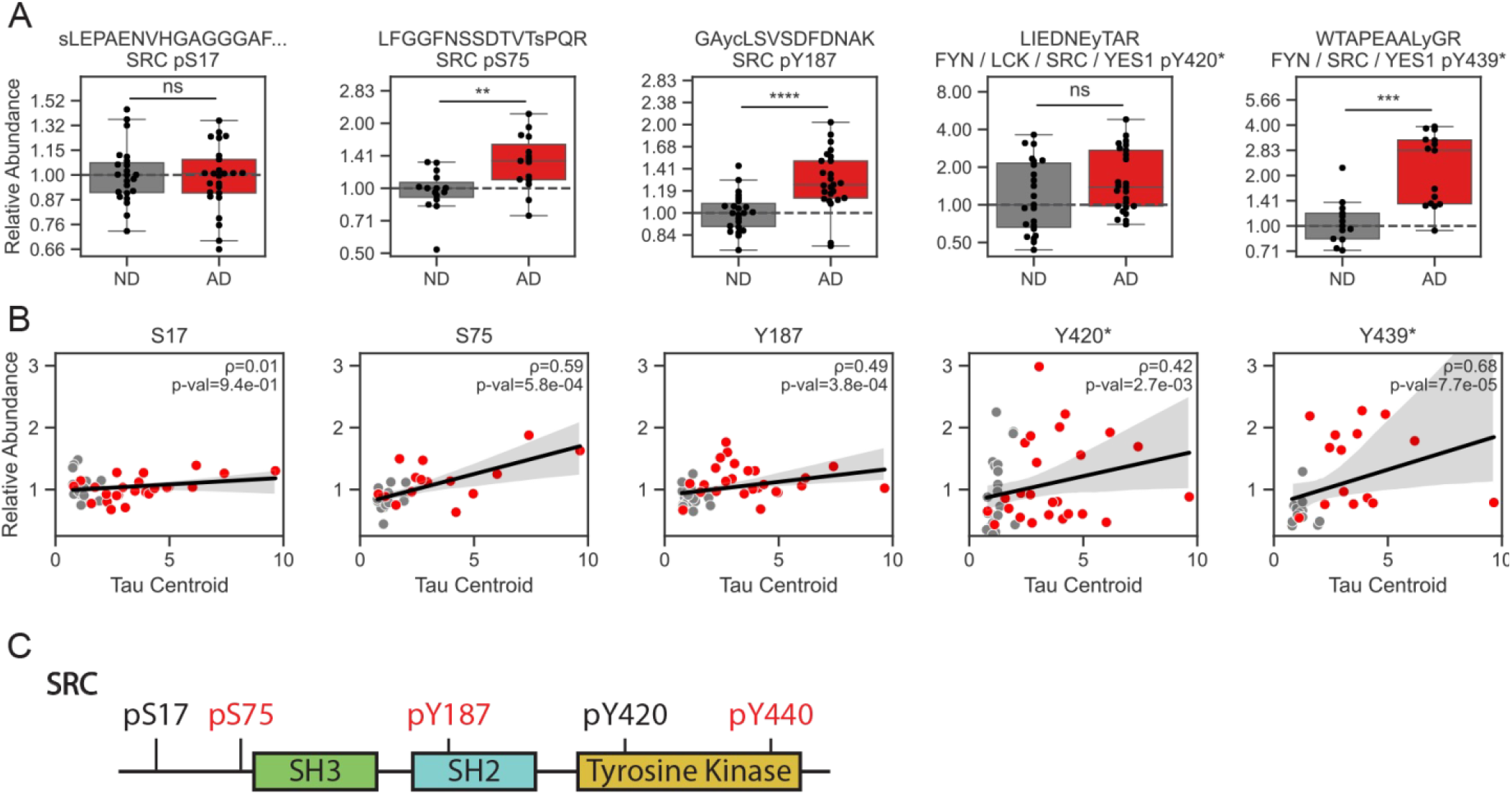
(A) Relative abundance of Src phosphorylation sites on pSer15, pSer75, pTyr187, pTyr420*, and pTyr439* in middle temporal gyrus (MTG) cortical brain tissue from patients with late-onset AD and age-matched, non-diseased (ND) controls. Phosphopeptides were identified using LC-MS/MS analysis and quantified using TMT10-plex. (B) Tau cluster centroid, containing the average of several Tau phosphopeptides, compared with Src phosphorylation levels across AD and ND samples. Spearman’s correlation coefficient and p-value are indicated in each panel. (C) Diagram illustrating Src domains and identified phosphorylation sites. AD or Tau-associated sites are colored red. **p < 1e-2; ***p < 1e-3; ****p < 1e-4.

### *Src* is upregulated in excitatory AD neurons

The emergence of large-scale single nuclei RNAseq (snRNAseq) data generated from post-mortem brain tissue from subjects affected by AD has offered new approaches for characterizing AD-associated molecular networks in a cell-specific manner. We recently generated snRNAseq superior frontal gyrus (SFG) profiles from 101 exceptionally well characterized subjects^50^, and interrogated this data to examine AD-associated Src mRNA expression in a cell-type specific manner. Consistent with our histochemical and protein quantification analyses, we observed that *Src* is significantly upregulated in excitatory neurons (Ex). Notably, *Src* is most highly expressed in excitatory neurons, both in AD and non-demented control (NC) samples, while inhibitory neurons (In) also display notable levels of Src expression. Importantly, the upregulation of Src in AD extends beyond excitatory neurons to include inhibitory neurons, emphasizing its broader involvement in neuronal populations within the context of AD. Examination of multiscale embedded gene co-expression networks^30^ constructed from Ex neurons identified transcription factor *NEUROD2* as a hub of the subnetwork that contains *Src*, and potential regulator of *Src* activity. We also observed positive correlations between *NEUROD2* and *Src* expression in at least five subtypes of excitatory neurons from another large-scale snRNA-seq profiling from postmortem brain tissues of AD clinical cohort^33^ (**Supplementary Table 4**). Visual inspection of photomicrographs confirmed the upregulation of *NEUROD2* in the SFG of AD patients compared to control. The observed positive correlation between *Src* and *NEUROD2* in excitatory neurons, along with the significant upregulation of *NEUROD2* in AD patients, raises the intriguing possibility that the observed increase in *Src* activity in AD is driven by transcription factor *NEUROD2*. Additional genetic perturbation studies of AD model systems to delineate this relationship may be warranted.

### Multiple Src phosphorylation sites are upregulated in AD

We next examined whether Src had altered kinase activity in AD by measuring its phosphorylation sites using mass spectrometry. We analyzed Src phosphorylation in MTG cortical brain (BA21) tissue from 25 AD and 23 non-demented (ND) cases using tandem-mass tag (TMT) 10-plex quantification, phosphotyrosine and global phosphoproteome enrichment, and untargeted LC-MS^2^ analysis^51^. This analysis quantified phosphopeptide mapping to Src pSer17, pSer75, pTyr187, pTyr420*, and pTyr439* across at least 28 patients (* phosphopeptide ambiguously mapped to multiple Src Family Kinases (SFKs)). Of these phosphorylation sites, Src pSer75, pTyr187, and pTyr439* were associated with AD (**Figure 5A**) or Tau phosphorylation levels (**Figure 5B**) which were also captured in this phosphoproteome analysis. Among these phosphorylation sites, pSer75 is a putative CDK5 substrate^52.^, pTyr187 is located on Src’s SH2 domain (**Figure 5C**) and shares homology with other SFK autoinhibitory sites^53^. Although the effects of pTyr439 are currently unknown, it is located on Src’s tyrosine kinase domain near pTyr420 which is crucial for kinase activity^54.^. The data shows that Src kinase undergoes distinct phosphorylation alterations in AD. These specific changes in Src kinase phosphorylation may provide a crucial molecular link for connecting Src kinase activity with AD pathogenesis.

**Figure 5:**
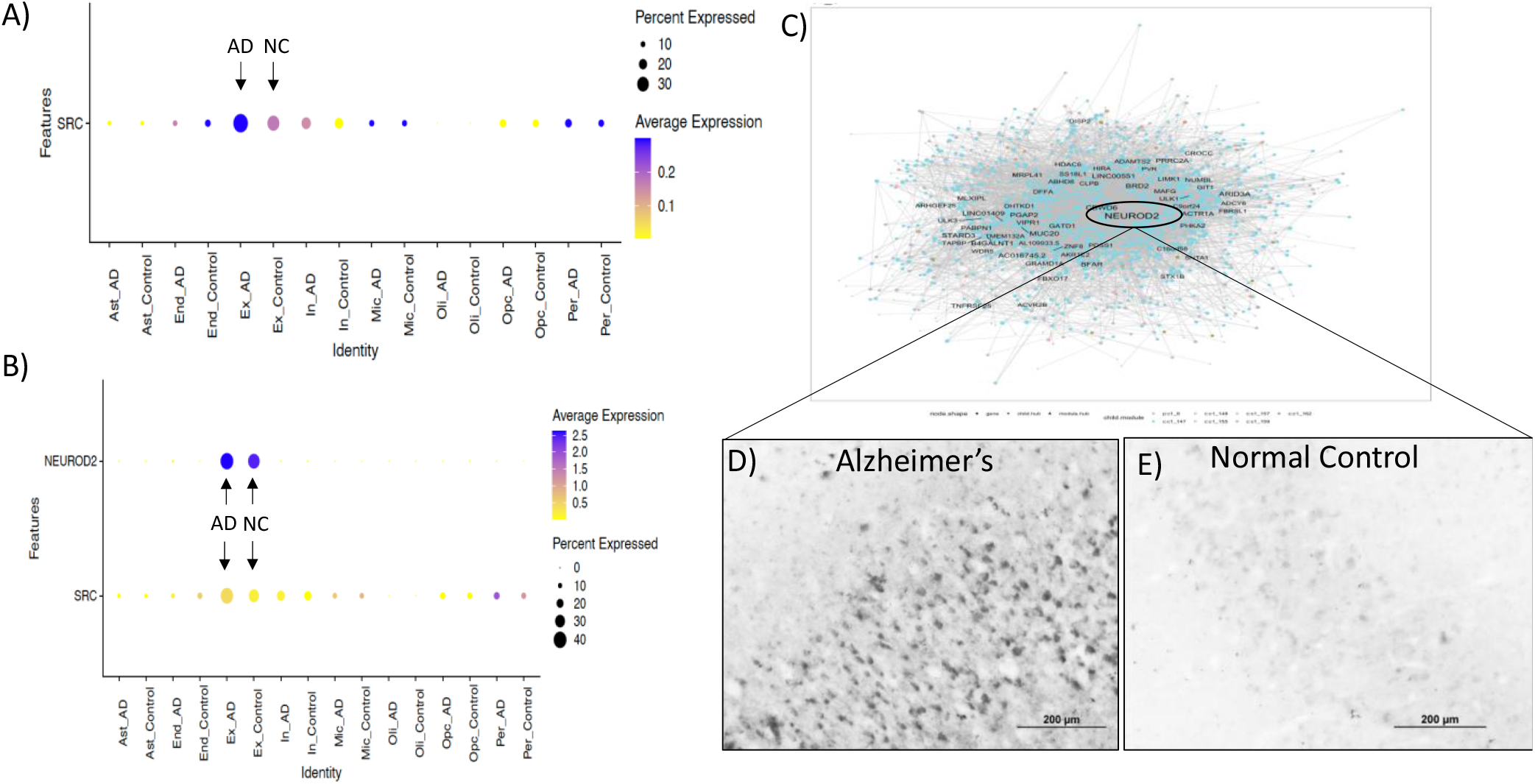
Src expression in Single nuclei RNAseq of 101 subjects stratified by cell type. A) Dot plot of cell types show that the most significant expression for Src is in excitatory neurons (Ex) in both AD and NC samples, followed by inhibitory neurons (In). AD Ex and In neurons were both significantly up-regulated compared to control (p=1.92E-115, and p=0.0005 respectively). All other cell types show weak Src expression levels. Multiscale Embedded Gene Co-expression Network Analysis identified the module hub NEUROD2 (B & C). A significant positive correlation is observed with Src and co-expression hub NEUROD2 in Ex neurons in both AD and NC samples. Representative photomicrographs show the up-regulation of NEUROD2 in AD SFG (D) compared to matched control (E).

### Tau and Src simulations

We next examined the implications of structural and interaction changes in Tau and Src through simulating Src-Tau interaction under 2 different conditions: (1) a microtubule (MT) with fragments of Tau bound (Figure 6A); and (2) the protofilaments of 2 Tau disease aggregates (Figure 6A), both of which are found in chronic traumatic encephalopathy (CTE) and AD^41,55^, termed Tau-CTE type 1 and Tau-CTE type 2. As Tau is known to be a stabilizing protein for MT, the first condition serves as an attempt to simulate Tau under a non-disease condition, with the second condition representing a disease scenario. Due to the disordered nature of Tau, a substantial amount of Tau residues is not resolved in structural models simulated here. However, among all the residues that are present (residues 252 to 379), they are all able to interact with Src through making contacts with the protein (Figure 6B); with those that are more solvent exposed interact with Src more frequently than those that are less exposed (Figure 6C). Further analyses show that Src residues that make contacts with Tau are different between the non-disease condition and the disease condition. For the non-disease condition, Tau-Src interaction is primarily observed for the first 110 Src residues in the Src structure employed (Figure 6D); for the disease conditions, Tau-Src interaction is primarily observed for residues 202 to residue 432 in the Src structure employed (Figure 6D). A less prominent difference is also observed between Tau-Src interaction and Src-MT interaction, where the set of Src residues that contact Tau with a normalized Src contact frequency larger than 0.4 is grossly different from that for Src-MT contacts (Figure 6D). Src residues that most frequently contact Tau or MT in our simulations are visualized in Figure 6E, where their spatial distributions were visually compared to the electrostatic profile of Src, suggesting that Src-Tau contact may prefer to occur at the electronegative region of Src while Src-MT contact may prefer an electropositive region of Src (Figure 6E).

**Figure 6:**
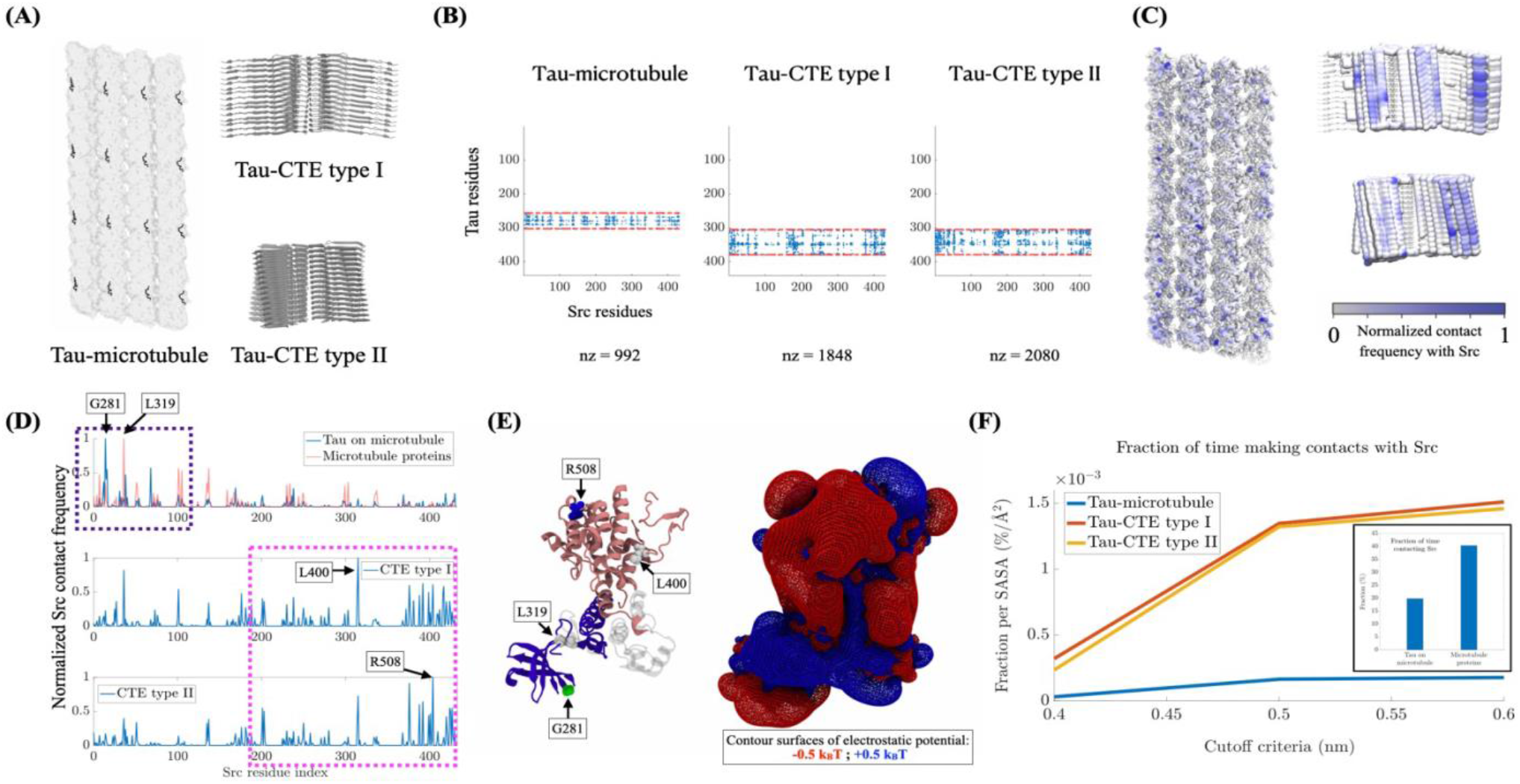
(A) Molecular images of a microtubule (MT) with fragments of Tau associated on it, and the protofilaments of 2 Tau disease aggregates, corresponding to CTE-type I & II. In these structural models, Tau residues are highlighted in gray. MT proteins are represented by a white, transparent protein scaffold. (B) Contact maps between Tau residues and Src residues computed from simulations. For each structural model employed, the minimal and maximal residue indices for Tau residues that are present are indicated by red, dashed lines. In-between these 2 lines are Tau residues that are available from the structure. Tau residues beyond those sandwiched by these 2 lines are unavailable. (C) Spatial distributions of Src associations hotspots on MT with Tau fragments and Tau disease aggregates. Each residue is colored according to its contact frequency with Src, normalized to have 1 being the maximal value. For clarity, residues with a normalized contact frequency with Src below 0.01 are all colored white. (D) Contact profile with Src. The frequency between each Src residue and each Tau or MT residue was measured and was normalized to have 1 being the maximal value for each structural model. Regions that capture significantly more Src-Tau contacts in the disease models than in the MT-Tau model are highlighted by a pink box, the region showing the opposite observation is highlighted by a blue box. Src residues that make most contact with either Tau (G281, L400, or R508) or MT (L319) in each structural model are highlighted by their residue IDs. (E) Molecular visualizations of key Src residues and Src electrostatic profiles. The pink region corresponds to residues highlighted by the pink box in (D). The same applies to the blue region. Key residues are colored according to their residue types (Blue: basic; Green: polar; White: hydrophobic). (F) Numbers of Src associations to either MT-Tau model or Tau disease models. The number of associations was measured in terms of the fraction of simulation time. For instance, in our current setting, a simulation of 2μs gives 200 trajectory frames. A value of 20% means observing Src associations for 200 × 20% = 40 frames. The comparison between the disease models and the MT-Tau models was made through the fraction per Solvent-Accessible Surface Area (SASA) so that the different geometrical shapes and sizes for our structural models can be accounted for.

Our simulations also show that Src associates with Tau much stronger than MT. For the non-disease condition, Src was observed to associate with either Tau or MT around 60% of the simulation time, among which, 20% was primarily associated with Tau and 40% was primarily associated with MT (Figure 6F). Src was classified to be primarily associated with Tau if Src makes more contacts with Tau than with MT. However, in the structural model for the non-disease condition, there are only 196 Tau residues, in contrast to the 20736 MT residues that are present. Thus, compared with MT, Tau, with only 196/20736 ≈ 0.095 (less than 1%) fraction of residues present, functions as the primarily association target for Src for 1/3 of the Src-Tau/MT associations observed. This strong affinity between Tau and Src is further demonstrated by comparing the non-disease condition to the disease condition with respect to the fraction of simulation time that Src is in contact with its association targets, namely MT with Tau for the non-disease condition and Tau aggregates for the disease condition. Our simulations show that Src contacts its association target much more frequently in the diseases condition than in the non-disease condition, up to 7-fold (Figure 6F). This upregulation shows differences between the 2 different structural models used for the disease condition, suggesting that the absence of MT to be the key factor underlying the upregulation while the fold states of Tau in the protofilaments contribute a less significant role in boosting Src-Tau associations.

## Discussion

The AD continuum is composed of multiple pathological features, most of which begin decades before clinical manifestations. The identification of molecular targets capable of modifying the earliest stages of AD prior to the onset of clinical symptoms is a critical pillar in the development of effective therapies for this devastating disease. Using publicly available and novel transcriptomic and proteomic data we have identified a disease-specific upregulation of the proto-oncogene Src kinase across multiple AD brain regions. Upregulation of Src is most strongly associated with early Tau marker MC1, and highly expressed in AD excitatory neurons. Brownian dynamics simulations of Src and Tau show that Tau aggregation induces Src contact potential 7-fold. *Src* phosphorylation by mass spectrometry identified increased phosphorylation on Ser75, Tyr187, and Tyr440 in AD. Together these results show that Src kinase is frequently active in AD neurons and may represent a therapeutic target for further evaluation.

### Importance of analyzing multiple brain regions

Alzheimer’s disease does not affect a single brain region. In fact, a pathological diagnosis of AD requires neurofibrillary tangle and amyloid plaque pathology in both the limbic and neocortical brain regions^47^. Most animal and human research performed in the dementia field emphasize pathological changes in the hippocampus, however, since this region is devastated at Braak stages IV and above, and removal of this brain region alone does not cause dementia but rather amnesia^56^, focusing solely on this region may limit our ability to disentangle the effects of established disease from its earliest drivers. To minimize these limitations, our approach was to analyze Src expression in multiple brain regions: hippocampus (BA27), middle temporal gyrus (BA21), entorhinal cortex (BA28), superior frontal gyrus (BA11) and in a comparatively unaffected brain region, the substantia nigra. Our results show that the most significantly impacted brain region with respect to Src expression was the entorhinal cortex (EC) followed by the hippocampus and neocortical subregions. Nigral Src immunoreactivity was minimally present in AD subjects and unreactive in control or PD subjects indicating that alterations in Src are AD-specific. It is not uncommon for AD subjects to show NFT pathology in basal ganglion structures^57^, but what was unexpected was the amount of neuropil immunoreactivity in the EC compared to other brain regions. Since the neuropil is comprised of mostly dendrites, axons, and synapses^44^, it provides a proxy measure of total neuronal connectivity within a tissue. The EC interconnects heavily with the neocortex and hippocampus, and is one of the earliest, if not the earliest brain region affected in AD^45^, indicating that dysregulation of Src occur in the EC long before other less reactive brain regions. The EC is the major input and output structure of the hippocampal formation, forming the nodal point in cortico-hippocampal circuits via layers II and III neurons^58^. Given the ECs importance in the cortical-hippocampal loop and memory consolidation, it will be important to determine if the same pattern of disease spreading associated with Tau (e.g., Braak staging) is also seen with Src. This may provide some insight into how early in the course of disease Src is up-regulated, and if the upregulation of Src predates Tau phosphorylation or if the two coincide.

### Co-occurrence with Tau markers

The trans-synaptic spread of Tau pathology in the CNS and its more recent systemic involvement has been an area of intense investigation. As such, systemic Tau has been shown to be a reliable biomarker of disease progression to aid in clinical assessments^59^. The biological tools used to identify Tau biomarkers in the periphery are largely predicated on disease-specific phospho-Tau epitopes. A recent study using p-tau231 indicated its early involvement in AD development^60^, and in our study we show a strong association with Src and p-tau231 (probability of p-tau231 given Src 61%,), but we show an even stronger association with the MC1 epitope (probability of MC1 given Src =100%), one of the earliest known disease-associated changes in Tau. In this conformationally dependent epitope the N-terminus of Tau interacts with its C-terminal third microtubule-binding repeat^61^. This MC1 conformation was determined to be highly soluble compared to the PHF1 epitope which is highly insoluble. Tau becomes insoluble after assembling into PHFs, suggesting that the change in Tau conformation from a relatively disordered soluble protein to the early conformationally altered MC1-Tau and the highly ordered PHF structure is crucial in the aggregation cascade. Importantly, the level of MC1 reactivity was shown to correlate with the severity and progression of AD with no reactivity against control brain extracts^46^.

The fact that the earliest disease states (Braak stage IV) were highly reactive for Src and concurrently reactive for the earliest Tau disease state (MC1) indicates a compelling association between Src activation and the early conformational changes in Tau, suggesting a potential pivotal role for this molecular interplay in the initiation and progression of tangle formation.

### *Src* Kinase in Neuronal Populations

The differential expression of *Src* kinase in distinct neuronal populations emerges as a compelling and thought-provoking dimension of this study. This observation invites a deeper exploration of the specific roles played by different types of neurons in the pathogenesis of AD and the potential involvement of *Src* kinase in these processes.

Notably, the heightened expression of *Src* kinase in AD excitatory neurons is particularly noteworthy. Excitatory neurons have been found to be more susceptible to pathological Tau accumulation in AD^62^. Augmented *Src* expression within excitatory neurons hints at their potential vulnerability to the influence of *Src* kinase within the context of AD. Multiscale co-expression networks constructed from excitatory neuronal snRNAseq implicate a potential regulatory relationship between transcription factor *NEUROD2* and *Src* kinase. These interactions may indicate critical roles in the modulation of responses to AD-related stressors and the development of Tau pathology, emphasizing the need for further investigations into the precise functional consequences of *Src* kinase upregulation in both excitatory and inhibitory neurons, including interactions with *NEUROD2*. This differential expression of *Src* kinase in neuronal populations carries significant implications for therapeutic strategies targeting this kinase. The prospect of selectively modulating *Src* kinase activity in highly expressed excitatory neurons offers a more targeted approach to mitigating Tau pathology and synaptic dysfunction in AD. Such selective modulation could potentially reduce side effects associated with broad kinase inhibition. Furthermore, the consideration of neuronal populations within intricate neuronal networks underscores the necessity of comprehending how *Src* kinase influences network-level functions in the context of AD. This understanding is crucial for elucidating the broader impact of the disease on brain function and for developing interventions that can address these intricate processes at multiple levels.

### Phosphorylation status of Src kinase in AD and its potential implications

Examining the phosphorylation status of Src kinase in AD unveils critical insights into the molecular changes associated with the disease. Our data reveals that Src kinase exhibits altered phosphorylation patterns in AD. These phosphorylation sites include pSer17, pSer75, pTyr187, pTyr420*, and pTyr439*. Among these phosphorylation sites, pSer75, pTyr187, and pTyr439* are associated with AD, suggesting that these specific phosphorylation events on Src are linked to AD. Additionally, the data indicates a correlation between the phosphorylation levels of Src at these sites and Tau phosphorylation levels, which is a significant finding in the context of AD. The data also provides insights into the potential functional implications of these phosphorylation changes. For instance, pSer75 is suggested to be a substrate for CDK5^52^, a kinase associated with neurodegeneration.^63^ pTyr187 is in the Src SH2 domain and shares homology with other sites involved in autoinhibition, suggesting a role in regulating Src activity. The impact of pTyr439 is currently unknown, but its location on the tyrosine kinase domain near pTyr420, which is crucial for kinase activity, implies potential relevance to Src function. This data suggests that changes in Src phosphorylation may be involved in the disease’s pathogenesis and, potentially, the phosphorylation of Tau. This provides a molecular link between Src and AD, strengthening the argument for its involvement in disease mechanisms. These data open up several future perspectives for research, by highlighting the need to explore the functional consequences of these specific phosphorylation events on Src in AD, particularly with regard to its role in Tau phosphorylation and the overall progression of the disease. Further investigations could focus on developing therapeutic strategies targeting Src in a phosphorylation site-specific manner to potentially mitigate AD-related pathology, providing a new avenue for treatment and intervention.

### Structural insights into Tau filaments and their interactions with Src

Detailed cryo-EM structures of Tau filaments from AD brains have deepened our understanding of the pathological changes associated with AD^55^. These findings underscore the importance of structural studies in deciphering the molecular basis of neurodegenerative diseases. The recognition of Paired Helical Filaments (PHFs) and Straight Filaments (SFs) as distinct structural polymorphs raises questions about their specific roles in AD progression. The observed interactions of specific residues of Tau with Src, particularly in association with AD, highlights a potential link between Tau aggregation and intracellular signaling pathways. Understanding the consequences of this interaction, such as how it affects neuronal function or contributes to AD pathogenesis, is an essential avenue for further investigation. Investigating the temporal relationship between Tau filament formation and the Tau-Src interaction, as well as the broader cellular and molecular consequences, will be vital. Moreover, understanding how structural variations in Tau filaments relate to disease severity and the clinical manifestation of AD is a promising avenue for further inquiry. The combined insights from the Tau filament structures and the Tau-Src interactions offer potential therapeutic implications. Targeting Tau filament formation or its interactions with Src could represent novel approaches for the development of mechanism-based therapies for AD and related neurodegenerative conditions. These findings provide a structural basis for future drug development efforts.

## Conclusion

The combined insights from this study not only enhance our understanding of the interaction of Src and Tau but also provide a foundation for future drug development efforts. Targeting Tau filament formation or its interactions with Src holds promise as a novel approach to developing mechanism-based therapies for AD and related neurodegenerative conditions. This is particularly significant given the availability of Src inhibitors, which could be repurposed or further developed for clinical trials in AD, potentially expediting the drug development process. This work underscores the importance of a multifaceted approach to decipher the complexity of AD and offers valuable perspectives for future research and potential treatments.

## Work Cited

1. Høgh, P. (2017). [Alzheimer’s disease]. Ugeskr Laeger 179.

2. Lane, C.A., Hardy, J., and Schott, J.M. (2018). Alzheimer’s disease. Eur J Neurol 25, 59–70. 10.1111/ene.13439.

3. Association, A.s. (2018). 2018 Alzheimer’s disease facts and figures. Alzheimer’s & Dementia 14, 367–429.

4. Gonneaud, J., Arenaza-Urquijo, E.M., Mezenge, F., Landeau, B., Gaubert, M., Bejanin, A., de Flores, R., Wirth, M., Tomadesso, C., Poisnel, G., et al. (2017). Increased florbetapir binding in the temporal neocortex from age 20 to 60 years. Neurology 89, 2438–2446. 10.1212/WNL.0000000000004733.

5. Serrano-Pozo, A., Frosch, M.P., Masliah, E., and Hyman, B.T. (2011). Neuropathological alterations in Alzheimer disease. Cold Spring Harbor perspectives in medicine 1, a006189.

6. Parsons, S.J., and Parsons, J.T. (2004). Src family kinases, key regulators of signal transduction. Oncogene 23, 7906–7909.

7. Cance, W.G., Craven, R.J., Bergman, M., Xu, L., Alitalo, K., and Liu, E.T. (1994). Rak, a novel nuclear tyrosine kinase expressed in epithelial cells. Cell growth & differentiation 5, 1347–1356.

8. Lee, J., Wang, Z., Luoh, S.-M., Wood, W.I., and Scadden, D.T. (1994). Cloning of FRK, a novel human intracellular SRC-like tyrosine kinaseencoding gene. Gene 138, 247–251.

9. Thuveson, M., Albrecht, D., Zurcher, G., Andres, A.-C., and Ziemiecki, A. (1995). Iyk, a novel intracellular protein tyrosine kinase differentially expressed in the mouse mammary gland and intestine. Biochemical and biophysical research communications 209, 582–589.

10. Amanchy, R., Zhong, J., Hong, R., Kim, J.H., Gucek, M., Cole, R.N., Molina, H., and Pandey, A. (2009). Identification of c-Src tyrosine kinase substrates in platelet-derived growth factor receptor signaling. Molecular oncology 3, 439–450.

11. Amanchy, R., Zhong, J., Molina, H., Chaerkady, R., Iwahori, A., Kalume, D.E., Grønborg, M., Joore, J., Cope, L., and Pandey, A. (2008). Identification of c-Src tyrosine kinase substrates using mass spectrometry and peptide microarrays. Journal of proteome research 7, 3900–3910.

12. Luo, W., Slebos, R.J., Hill, S., Li, M., Brábek, J., Amanchy, R., Chaerkady, R., Pandey, A., Ham, A.-J.L., and Hanks, S.K. (2008). Global impact of oncogenic Src on a phosphotyrosine proteome. Journal of proteome research 7, 3447–3460.

13. Arbesú, M., Maffei, M., Cordeiro, T.N., Teixeira, J.M., Pérez, Y., Bernadó, P., Roche, S., and Pons, M. (2017). The unique domain forms a fuzzy intramolecular complex in Src family kinases. Structure 25, 630–640. e634.

14. Combs, C.K., Bates, P., Karlo, J.C., and Landreth, G.E. (2001). Regulation of beta-amyloid stimulated proinflammatory responses by peroxisome proliferator-activated receptor alpha. Neurochem Int 39, 449–457. 10.1016/s0197-0186(01)00052-3.

15. Combs, C.K., Johnson, D.E., Cannady, S.B., Lehman, T.M., and Landreth, G.E. (1999). Identification of microglial signal transduction pathways mediating a neurotoxic response to amyloidogenic fragments of beta-amyloid and prion proteins. J Neurosci 19, 928–939. 10.1523/jneurosci.19-03-00928.1999.

16. Bamberger, M.E., Harris, M.E., McDonald, D.R., Husemann, J., and Landreth, G.E. (2003). A cell surface receptor complex for fibrillar beta-amyloid mediates microglial activation. J Neurosci 23, 2665–2674. 10.1523/jneurosci.23-07-02665.2003.

17. Combs, C.K., Karlo, J.C., Kao, S.C., and Landreth, G.E. (2001). beta-Amyloid stimulation of microglia and monocytes results in TNFalpha-dependent expression of inducible nitric oxide synthase and neuronal apoptosis. J Neurosci 21, 1179–1188. 10.1523/jneurosci.21-04-01179.2001.

18. McDonald, D.R., Bamberger, M.E., Combs, C.K., and Landreth, G.E. (1998). beta-Amyloid fibrils activate parallel mitogen-activated protein kinase pathways in microglia and THP1 monocytes. J Neurosci 18, 4451–4460. 10.1523/jneurosci.18-12-04451.1998.

19. Um, J.W., Nygaard, H.B., Heiss, J.K., Kostylev, M.A., Stagi, M., Vortmeyer, A., Wisniewski, T., Gunther, E.C., and Strittmatter, S.M. (2012). Alzheimer amyloid-β oligomer bound to postsynaptic prion protein activates Fyn to impair neurons. Nature neuroscience 15, 1227–1235.

20. Larson, M., Sherman, M.A., Amar, F., Nuvolone, M., Schneider, J.A., Bennett, D.A., Aguzzi, A., and Lesné, S.E. (2012). The complex PrPc-Fyn couples human oligomeric Aβ with pathological Tau changes in Alzheimer’s disease. Journal of Neuroscience 32, 16857–16871.

21. Haas, L.T., Salazar, S.V., Kostylev, M.A., Um, J.W., Kaufman, A.C., and Strittmatter, S.M. (2016). Metabotropic glutamate receptor 5 couples cellular prion protein to intracellular signalling in Alzheimer’s disease. Brain 139, 526–546.

22. Um, J.W., Kaufman, A.C., Kostylev, M., Heiss, J.K., Stagi, M., Takahashi, H., Kerrisk, M.E., Vortmeyer, A., Wisniewski, T., and Koleske, A.J. (2013). Metabotropic glutamate receptor 5 is a coreceptor for Alzheimer aβ oligomer bound to cellular prion protein. Neuron 79, 887–902.

23. Polonio-Vallon, T., Kirkpatrick, J., Krijgsveld, J., and Hofmann, T.G. (2014). Src kinase modulates the apoptotic p53 pathway by altering HIPK2 localization. Cell Cycle 13, 115–125. 10.4161/cc.26857.

24. Beach, T.G., Adler, C.H., Sue, L.I., Serrano, G., Shill, H.A., Walker, D.G., Lue, L., Roher, A.E., Dugger, B.N., Maarouf, C., et al. (2015). Arizona Study of Aging and Neurodegenerative Disorders and Brain and Body Donation Program. Neuropathology 35, 354–389. 10.1111/neup.12189.

25. Braak, H., Alafuzoff, I., Arzberger, T., Kretzschmar, H., and Del Tredici, K. (2006). Staging of Alzheimer disease-associated neurofibrillary pathology using paraffin sections and immunocytochemistry. Acta Neuropathol 112, 389–404. 10.1007/s00401-006-0127-z.

26. Mastroeni, D., McKee, A., Grover, A., Rogers, J., and Coleman, P.D. (2009). Epigenetic differences in cortical neurons from a pair of monozygotic twins discordant for Alzheimer’s disease. PLoS One 4, e6617. 10.1371/journal.pone.0006617.

27. Mastroeni, D., Delvaux, E., Nolz, J., Tan, Y., Grover, A., Oddo, S., and Coleman, P.D. (2015). Aberrant intracellular localization of H3k4me3 demonstrates an early epigenetic phenomenon in Alzheimer’s disease. Neurobiol Aging 36, 3121–3129. 10.1016/j.neurobiolaging.2015.08.017.

28. Mastroeni, D., Chouliaras, L., Grover, A., Liang, W.S., Hauns, K., Rogers, J., and Coleman, P.D. (2013). Reduced RAN expression and disrupted transport between cytoplasm and nucleus; a key event in Alzheimer’s disease pathophysiology. PLoS One 8, e53349. 10.1371/journal.pone.0053349.

29. Wang, Q., Antone, J., Alsop, E., Reiman, R., Funk, C., Bendl, J., Dudley, J.T., Liang, W.S., Karr, T.L., Roussos, P., et al. (2023). A public resource of single cell transcriptomes and multiscale networks from persons with and without Alzheimer’s disease. bioRxiv. 10.1101/2023.10.20.563319.

30. Song, W.M., and Zhang, B. (2015). Multiscale Embedded Gene Co-expression Network Analysis. PLoS Comput Biol 11, e1004574. 10.1371/journal.pcbi.1004574.

31. Hao, Y., Hao, S., Andersen-Nissen, E., Mauck, W.M., 3rd, Zheng, S., Butler, A., Lee, M.J., Wilk, A.J., Darby, C., Zager, M., et al. (2021). Integrated analysis of multimodal single-cell data. Cell 184, 3573–3587 e3529. 10.1016/j.cell.2021.04.048.

32. Finak, G., McDavid, A., Yajima, M., Deng, J., Gersuk, V., Shalek, A.K., Slichter, C.K., Miller, H.W., McElrath, M.J., Prlic, M., et al. (2015). MAST: a flexible statistical framework for assessing transcriptional changes and characterizing heterogeneity in single-cell RNA sequencing data. Genome Biol 16, 278. 10.1186/s13059-015-0844-5.

33. Mathys, H., Peng, Z., Boix, C.A., Victor, M.B., Leary, N., Babu, S., Abdelhady, G., Jiang, X., Ng, A.P., Ghafari, K., et al. (2023). Single-cell atlas reveals correlates of high cognitive function, dementia, and resilience to Alzheimer’s disease pathology. Cell 186, 4365–4385.e4327. 10.1016/j.cell.2023.08.039.

34. Ritchie, M.E., Phipson, B., Wu, D., Hu, Y., Law, C.W., Shi, W., and Smyth, G.K. (2015). limma powers differential expression analyses for RNA-sequencing and microarray studies. Nucleic Acids Res 43, e47. 10.1093/nar/gkv007.

35. Morshed, N., Lee, M.J., Rodriguez, F.H., Lauffenburger, D.A., Mastroeni, D., and White, F.M. (2021). Quantitative phosphoproteomics uncovers dysregulated kinase networks in Alzheimer’s disease. Nat Aging 1, 550–565. 10.1038/s43587-021-00071-1.

36. Morshed, N., Ralvenius, W.T., Nott, A., Watson, L.A., Rodriguez, F.H., Akay, L.A., Joughin, B.A., Pao, P.C., Penney, J., LaRocque, L., et al. (2020). Phosphoproteomics identifies microglial Siglec-F inflammatory response during neurodegeneration. Mol Syst Biol 16, e9819. 10.15252/msb.20209819.

37. Breitenlechner, C.B., Kairies, N.A., Honold, K., Scheiblich, S., Koll, H., Greiter, E., Koch, S., Schafer, W., Huber, R., and Engh, R.A. (2005). Crystal structures of active SRC kinase domain complexes. J Mol Biol 353, 222–231. 10.1016/j.jmb.2005.08.023.

38. Berman, H.M., Westbrook, J., Feng, Z., Gilliland, G., Bhat, T.N., Weissig, H., Shindyalov, I.N., and Bourne, P.E. (2000). The Protein Data Bank. Nucleic Acids Res 28, 235–242. 10.1093/nar/28.1.235.

39. Falcon, B., Zivanov, J., Zhang, W., Murzin, A.G., Garringer, H.J., Vidal, R., Crowther, R.A., Newell, K.L., Ghetti, B., Goedert, M., and Scheres, S.H.W. (2019). Novel tau filament fold in chronic traumatic encephalopathy encloses hydrophobic molecules. Nature 568, 420–423. 10.1038/s41586-019-1026-5.

40. Fitzpatrick, A.W.P., Falcon, B., He, S., Murzin, A.G., Murshudov, G., Garringer, H.J., Crowther, R.A., Ghetti, B., Goedert, M., and Scheres, S.H.W. (2017). Cryo-EM structures of tau filaments from Alzheimer’s disease. Nature 547, 185–190. 10.1038/nature23002.

41. Lovestam, S., Koh, F.A., van Knippenberg, B., Kotecha, A., Murzin, A.G., Goedert, M., and Scheres, S.H.W. (2022). Assembly of recombinant tau into filaments identical to those of Alzheimer’s disease and chronic traumatic encephalopathy. Elife 11. 10.7554/eLife.76494.

42. Chan, C.K., Singharoy, A., and Tajkhorshid, E. (2022). Anionic Lipids Confine Cytochrome c(2) to the Surface of Bioenergetic Membranes without Compromising Its Interaction with Redox Partners. Biochemistry 61, 385–397. 10.1021/acs.biochem.1c00696.

43. Zhang, X., Gao, F., Wang, D., Li, C., Fu, Y., He, W., and Zhang, J. (2018). Tau Pathology in Parkinson’s Disease. Front Neurol 9, 809. 10.3389/fneur.2018.00809.

44. Spocter, M.A., Hopkins, W.D., Barks, S.K., Bianchi, S., Hehmeyer, A.E., Anderson, S.M., Stimpson, C.D., Fobbs, A.J., Hof, P.R., and Sherwood, C.C. (2012). Neuropil distribution in the cerebral cortex differs between humans and chimpanzees. J Comp Neurol 520, 2917–2929. 10.1002/cne.23074.

45. Braak, H., and Braak, E. (1992). The human entorhinal cortex: normal morphology and lamina-specific pathology in various diseases. Neurosci Res 15, 6–31. 10.1016/0168-0102(92)90014-4.

46. Weaver, C.L., Espinoza, M., Kress, Y., and Davies, P. (2000). Conformational change as one of the earliest alterations of tau in Alzheimer’s disease. Neurobiol Aging 21, 719–727. 10.1016/s0197-4580(00)00157-3.

47. DeTure, M.A., and Dickson, D.W. (2019). The neuropathological diagnosis of Alzheimer’s disease. Mol Neurodegener 14, 32. 10.1186/s13024-019-0333-5.

48. Kerchner, G.A., Hess, C.P., Hammond-Rosenbluth, K.E., Xu, D., Rabinovici, G.D., Kelley, D.A., Vigneron, D.B., Nelson, S.J., and Miller, B.L. (2010). Hippocampal CA1 apical neuropil atrophy in mild Alzheimer disease visualized with 7-T MRI. Neurology 75, 1381–1387. 10.1212/WNL.0b013e3181f736a1.

49. Moloney, C.M., Lowe, V.J., and Murray, M.E. (2021). Visualization of neurofibrillary tangle maturity in Alzheimer’s disease: A clinicopathologic perspective for biomarker research. Alzheimers Dement 17, 1554–1574. 10.1002/alz.12321.

50. Wang, Q., Antone, J., Alsop, E., Reiman, R., Funk, C., Bendl, J., Dudley, J.T., Liang, W.S., Karr, T.L., Roussos, P., et al. (2023). A public resource of single cell transcriptomes and multiscale networks from persons with and without Alzheimer’s disease. bioRxiv, 2023.2010.2020.563319. 10.1101/2023.10.20.563319.

51. Nader Morshed, M.J.L., Felicia H. Rodriguez, Douglas A. Lauffenburger, Diego Mastroeni & Forest M. White (2021). Quantitative phosphoproteomics uncovers dysregulated kinase networks in Alzheimer’s disease. Nature Aging, 505–565. 10.1038/s43587-021-00071-1.

52. Pan, Q., Qiao, F., Gao, C., Norman, B., Optican, L., and Zelenka, P.S. (2011). Cdk5 targets active Src for ubiquitin-dependent degradation by phosphorylating Src(S75). Cell Mol Life Sci 68, 3425–3436. 10.1007/s00018-011-0638-1.

53. Weir, M.E., Mann, J.E., Corwin, T., Fulton, Z.W., Hao, J.M., Maniscalco, J.F., Kenney, M.C., Roman Roque, K.M., Chapdelaine, E.F., Stelzl, U., et al. (2016). Novel autophosphorylation sites of Src family kinases regulate kinase activity and SH2 domain-binding capacity. FEBS Lett 590, 1042–1052. 10.1002/1873-3468.12144.

54. Matrone, C., Petrillo, F., Nasso, R., and Ferretti, G. (2020). Fyn Tyrosine Kinase as Harmonizing Factor in Neuronal Functions and Dysfunctions. Int J Mol Sci 21. 10.3390/ijms21124444.

55. Tarutani, A., Lovestam, S., Zhang, X., Kotecha, A., Robinson, A.C., Mann, D.M.A., Saito, Y., Murayama, S., Tomita, T., Goedert, M., et al. (2023). Cryo-EM structures of tau filaments from SH-SY5Y cells seeded with brain extracts from cases of Alzheimer’s disease and corticobasal degeneration. FEBS Open Bio 13, 1394–1404. 10.1002/2211-5463.13657.

56. Scoville, W.B., and Milner, B. (2000). Loss of recent memory after bilateral hippocampal lesions. 1957. J Neuropsychiatry Clin Neurosci 12, 103–113. 10.1176/jnp.12.1.103.

57. Hamasaki, H., Honda, H., Suzuki, S.O., Shijo, M., Ohara, T., Hatabe, Y., Okamoto, T., Ninomiya, T., and Iwaki, T. (2019). Tauopathy in basal ganglia involvement is exacerbated in a subset of patients with Alzheimer’s disease: The Hisayama study. Alzheimers Dement (Amst) 11, 415–423. 10.1016/j.dadm.2019.04.008.

58. Nilssen, E.S., Doan, T.P., Nigro, M.J., Ohara, S., and Witter, M.P. (2019). Neurons and networks in the entorhinal cortex: A reappraisal of the lateral and medial entorhinal subdivisions mediating parallel cortical pathways. Hippocampus 29, 1238–1254. 10.1002/hipo.23145.

59. Arastoo, M., Lofthouse, R., Penny, L.K., Harrington, C.R., Porter, A., Wischik, C.M., and Palliyil, S. (2020). Current Progress and Future Directions for Tau-Based Fluid Biomarker Diagnostics in Alzheimer’s Disease. Int J Mol Sci 21. 10.3390/ijms21228673.

60. Ashton, N.J., Benedet, A.L., Pascoal, T.A., Karikari, T.K., Lantero-Rodriguez, J., Brum, W.S., Mathotaarachchi, S., Therriault, J., Savard, M., Chamoun, M., et al. (2022). Cerebrospinal fluid p-tau231 as an early indicator of emerging pathology in Alzheimer’s disease. EBioMedicine 76, 103836. 10.1016/j.ebiom.2022.103836.

61. Jicha, G.A., Bowser, R., Kazam, I.G., and Davies, P. (1997). Alz-50 and MC-1, a new monoclonal antibody raised to paired helical filaments, recognize conformational epitopes on recombinant tau. J Neurosci Res 48, 128–132. 10.1002/(sici)1097-4547(19970415)48:2<128::aid-jnr5>3.0.co;2-e.

62. Fu, H., Possenti, A., Freer, R., Nakano, Y., Hernandez Villegas, N.C., Tang, M., Cauhy, P.V.M., Lassus, B.A., Chen, S., Fowler, S.L., et al. (2019). A tau homeostasis signature is linked with the cellular and regional vulnerability of excitatory neurons to tau pathology. Nat Neurosci 22, 47–56. 10.1038/s41593-018-0298-7.

63. Liu, S.L., Wang, C., Jiang, T., Tan, L., Xing, A., and Yu, J.T. (2016). The Role of Cdk5 in Alzheimer’s Disease. Mol Neurobiol 53, 4328–4342. 10.1007/s12035-015-9369-x.

