## Supplemental Tables 1-4 for "Upregulation of the Proto-Oncogene Src Kinase in Alzheimer’s Disease: From Molecular Interactions to Therapeutic Potential"

#### Supplementary Table 1: Sample Demographics

Immunohistochemistry Data from 28 samples (Alzheimer's disease - AD = 10; Parkinson's disease - PD = 9; Nondemented or control - ND = 9) from 5 brain regions: hippocampus (BA27), superior frontal gyrus (BA11), middle temporal gyrus (BA21), entorhinal cortex (BA28) and substantia nigra pars compacta. There was no significant difference between sample age at time of death or post-mortem interval between AD, PD, and ND samples. Additionally, there was no significance difference between expired age, postmortem interval, ApoE status, or MMSE within or between AD, ND, or PD.

Western Blot Data from 12 samples (Alzheimer's disease - AD = 6; Nondemented or control - ND = 6) from the middle temporal gyrus (BA21). There was no significant difference between sample age at time of death or post-mortem interval between AD and ND samples. Additionally, there was no significance difference between expired age, postmortem interval, ApoE status, or MMSE within or between AD and ND.

| **Sample Metadata for Histochemical Data** | | | | |
|---|---|---|---|---|
| | **AD** | **PD** | **ND** | **Statistics** |
| ***n*** | 10 | 9 | 9 | |
| **Expired Age (years)** | 71.5 ± 8.21 | 72.3 ± 5.16 | 72.4 ± 6.12 | F = 0.1625*<br>p-value = 0.852 |
| **Sex** | 5 males/5 females | 5 males/4 females | 5 males/4 females | |
| **Post-mortem Interval (hours)** | 3.41 ± 0.99 | 2.79 ± 1.22 | 3.15 ± 0.98 | F = 1.1685*<br>p-value = 0.331 |
| **ApoE** | 2/3 = 0<br>3/3 =9<br>3/4 =1<br>4/4 = 0 | 2/3 = 1<br>3/3 =8<br>3/4 = 0<br>4/4 = 0 | 2/3 = 1<br>3/3 = 8<br>3/4 = 0<br>4/4 = 0 | |
| **Braak Stage** | I = 0 | I = 0 | I = 0 | |

| | | | |
|---|---|---|---|
| | II = 0<br>III = 0<br>IV = 10<br>V = 0<br>VI = 0 | II = 6<br>III = 3<br>IV = 0<br>V = 0<br>VI = 0 | II = 5<br>III = 4<br>IV = 0<br>V = 0<br>VI = 0 |
| **MMSE (last test score)** | 6.50 ± 8.74 | 21.50 ± 3.56 | 29.67 ± 0.58 |

| **Sample Metadata for Western Blot Data** | | | |
|---|---|---|---|
| | **AD** | **ND** | **Statistics** |
| ***n*** | 6 | 6 | |
| **Expired Age (years)** | 75.5 ± 1.41 | 75.2 ± 0.98 | p-value = 0.191 |
| **Sex** | 3 males/3 females | 3 males/3 females | |
| **Post-mortem Interval (hours)** | 2.91 ± 0.88 | 2.88 ± 1.22 | p-value = 0.875 |
| **ApoE** | 2/3 = 0<br>3/3 =6<br>3/4 =0<br>4/4 = 0 | 2/3 = 0<br>3/3 =6<br>3/4 = 0<br>4/4 = 0 | |
| **Braak Stage** | I = 0<br>II = 0<br>III = 0<br>IV = 6<br>V = 0<br>VI = 0 | I = 0<br>II = 4<br>III = 2<br>IV = 0<br>V = 0<br>VI = 0 | |
| **MMSE (last test score)** | 5.98 ± 9.44 | 29.01 ± 2.11 | |

### Supplementary Table 2: Primary & Secondary Antibodies/Sources/Dilutions

| | | Primary Antibodies | | | |
|---|---|---|---|---|---|
| Supplier | Protein Target | Catalogue # | Species | Clonality | Dilution |
| Abcam | NeuroD2 | ab104430 | Rabbit | Polyclonal | 1:500 |
| Abcam | Src | ab109381 | Rabbit | Monoclonal | 1:1000 (IHC) 1:5000 (WB) |
| Peter Davies | MC1 | N/A | Mouse | Monoclonal | 1:2000 |
| Peter Davies | PHF1 | N/A | Mouse | Monoclonal | 1:2000 |
| Peter Davies | Cp13 | N/A | Mouse | Monoclonal | 1:2000 |
| Peter Davies | Pg5 | N/A | Mouse | Monoclonal | 1:2000 |
| Thermo Fisher | pTau231 | 35-5200 | Mouse | Monoclonal | 1:1000 |
| | | Secondary Antibodies | | | |
| Supplier | Target | Catalogue # | Species | Clonality | |
| Thermo Fisher | Alexa Fluor 488, Anti-Rabbit IgG | A-11005 | Goat | Polyclonal | 1:2000 |
| Thermo Fisher | Alexa Fluor 594, Anti-Mouse IgG | A-11012 | Goat | Polyclonal | 1:2000 |
| Vector | Anti-Mouse IgG, biotinylated | BP-9200-50 | Goat | Polyclonal | 1:1000 |
| Vector | Anti-Rabbit IgG, biotinylated | BP-9100-50 | Goat | Polyclonal | 1:1000 |
| Thermo Fisher | Anti-Rabbit IgG, HRP | 31460 | Goat | Polyclonal | 1:5000 |
| | | Control | | | |
| Supplier | Target | Catalogue # | Species | Clonality | |
| Cell Signaling | Src Blocking Peptide | #1235 | N/A | N/A | 1:1 |
| Abcam | Beta Actin | ab8226 | Mouse | Monoclonal | 1:5000 |
| Invitrogen | DAPI | D1306 | N/A | N/A | 1:20000 |

### Supplementary Table 3: Probability of Src given Tau

| Src/MC1 | | | | | |
|---|---|---|---|---|---|
| Sample | total of neurons counted | MC1+ | Src+ | MC1+/SRC+ | Probability of Src given MC1 |
| sample 1 | 32 | 17 | 22 | 17 | 100% |
| sample 2 | 38 | 23 | 28 | 23 | 100% |
| sample 3 | 41 | 28 | 31 | 28 | 100% |
| sample 4 | 28 | 20 | 23 | 20 | 100% |
| sample 5 | 39 | 30 | 37 | 30 | 100% |
| sample 6 | 33 | 21 | 29 | 21 | 100% |
| sample 7 | 48 | 33 | 40 | 33 | 100% |
| sample 8 | 36 | 21 | 28 | 21 | 100% |
| sample 9 | 49 | 38 | 45 | 38 | 100% |
| sample 10 | 40 | 20 | 28 | 20 | |
| mean # of neurons | 38.4 | | | | 100% |
| mean probability | | | | | |

| Src/CP13 | | | | | |
|---|---|---|---|---|---|
| sample | total of neurons counted | CP13 reactive | Src reactive | CP13+/SRC+ | Probability of Src given CP13 |
| sample 1 | 39 | 17 | 25 | 10 | 59% |
| sample 2 | 48 | 19 | 32 | 10 | 53% |
| sample 3 | 38 | 18 | 27 | 9 | 50% |
| sample 4 | 33 | 14 | 23 | 10 | 71% |
| sample 5 | 38 | 20 | 29 | 14 | 70% |
| sample 6 | 37 | 21 | 29 | 10 | 48% |
| sample 7 | 40 | 22 | 36 | 14 | 64% |
| sample 8 | 38 | 12 | 22 | 7 | 58% |
| sample 9 | 44 | 24 | 40 | 16 | 67% |
| sample 10 | 38 | 14 | 31 | 10 | 71% |
| mean # of neurons | 39.3 | | | | |
| mean probability | | | | | 61% |

| Src/pTau213 | | | | | |
|---|---|---|---|---|---|
| sample | total of neurons counted | Tau 213 reacti | Src reactive | Tau213+/SRC | Probability of Src given pTau 213 |
| sample 1 | 39 | 12 | 28 | 8 | 67% |
| sample 2 | 32 | 19 | 28 | 10 | 53% |
| sample 3 | 40 | 20 | 34 | 10 | 50% |
| sample 4 | 31 | 13 | 28 | 9 | 69% |
| sample 5 | 44 | 23 | 37 | 14 | 61% |
| sample 6 | 29 | 12 | 29 | 7 | 58% |
| sample 7 | 40 | 19 | 31 | 11 | 58% |
| sample 8 | 31 | 11 | 29 | 7 | 64% |
| sample 9 | 44 | 22 | 40 | 12 | 55% |
| sample 10 | 31 | 20 | 28 | 10 | 50% |
| mean # of neurons | 36.1 | | | | |
| mean probability | | | | | 58% |

| Src/pg5 | | | | | |
|---|---|---|---|---|---|
| sample | total of neurons counted | PG5 reactive | Src reactive | pg5+/SRC+ | Probability of Src given pg5 |
| sample 1 | 36 | 17 | 22 | 9 | 53% |
| sample 2 | 31 | 23 | 28 | 12 | 52% |
| sample 3 | 39 | 28 | 31 | 13 | 46% |
| sample 4 | 33 | 20 | 23 | 12 | 60% |
| sample 5 | 48 | 30 | 37 | 15 | 50% |
| sample 6 | 27 | 21 | 21 | 11 | 52% |
| sample 7 | 39 | 33 | 31 | 21 | 64% |
| sample 8 | 41 | 31 | 38 | 18 | 58% |
| sample 9 | 38 | 19 | 21 | 8 | 42% |
| sample 10 | 41 | 20 | 27 | 12 | 60% |
| mean # of neurons | 37.3 | | | | |
| mean probability | | | | | 54% |

| Src/PHF1 | | | | | |
|---|---|---|---|---|---|
| sample | total of neurons counted | PHF1 reactive | Src reactive | PHF1+/SRC+ | Probability of Src given PHF1 |
| sample 1 | 43 | 9 | 21 | 4 | 44% |
| sample 2 | 33 | 12 | 24 | 3 | 25% |
| sample 3 | 44 | 20 | 29 | 10 | 50% |
| sample 4 | 38 | 13 | 19 | 6 | 46% |
| sample 5 | 37 | 18 | 21 | 6 | 33% |
| sample 6 | 31 | 12 | 21 | 4 | 33% |
| sample 7 | 35 | 10 | 31 | 4 | 40% |
| sample 8 | 39 | 21 | 31 | 12 | 57% |
| sample 9 | 44 | 18 | 40 | 8 | 44% |
| sample 10 | 37 | 17 | 26 | 9 | 53% |
| mean # of neurons | 38.1 | | | | |
| mean probability | | | | | 43% |

#### Supplementary Table 4: NEUROD2/SRC correlation in Ex subtypes.

| excitatory neuron subtype | beta correlat | p value | FDR | # of nuclei |
|---|---|---|---|---|
| Exc L2-3 CBLN2 LINC02306 | 0.0163 | 5.80E-01 | 6.77E-01 | 296,936 |
| Exc L3-4 RORB CUX2 | 0.0842 | 2.36E-04 | **3.30E-03** | 184,784 |
| Exc L3-5 RORB PLCH1 | 0.0652 | 4.71E-02 | 1.10E-01 | 37,949 |
| Exc L4-5 RORB GABRG1 | 0.0753 | 1.35E-02 | **3.78E-02** | 79,361 |
| Exc L4-5 RORB IL1RAPL2 | 0.0797 | 1.20E-03 | **6.39E-03** | 119,435 |
| Exc L5 ET | 0.1124 | 1.37E-03 | **6.39E-03** | 3,454 |
| Exc L5-6 RORB LINC02196 | 0.0365 | 3.04E-01 | 4.73E-01 | 22,343 |
| Exc L5/6 IT Car3 | -0.0004 | 9.93E-01 | 9.93E-01 | 18,371 |
| Exc L5/6 NP | 0.0522 | 2.64E-01 | 4.62E-01 | 17,247 |
| Exc L6 CT | 0.0688 | 1.41E-01 | 2.82E-01 | 23,073 |
| Exc L6 THEMIS NFIA | 0.0368 | 3.38E-01 | 4.73E-01 | 66,676 |
| Exc L6b | 0.0645 | 1.07E-02 | **3.75E-02** | 25,055 |
| Exc NRGN | 0.0174 | 7.64E-01 | 8.23E-01 | 45,859 |
| Exc RELN CHD7 | 0.0282 | 5.43E-01 | 6.77E-01 | 102,687 |
